# Emotional valence of conspecific vocalizations modulates auditory and limbic brain activity in juvenile pigs

**DOI:** 10.64898/2026.05.15.725583

**Authors:** Paul Coudert, Thibault Dussol, Yann Serrand, Nicolas Coquery, Stéphane Laurent, Hervé Saint-Jalmes, Gwenaelle Creff, Céline Tallet, Benoît Godey, David Val-Laillet, Pierre-Antoine Eliat

## Abstract

Pig vocalizations convey information about the emotional states of individuals, varying with arousal and valence. Studies show that different call types reflect distinct emotional contexts and social interactions for the receivers. However, little is known about the brain mechanisms behind the perception of conspecifics’ vocalizations. This study used BOLD fMRI to explore how pigs’ brains respond to emotionally varied vocalizations, with the aim to identify activity in regions linked to emotion, reward, and social processing.

Eight healthy 2-month-old pigs underwent auditory brainstem response (ABR) testing and BOLD fMRI to assess brain responses to pig vocalizations with different hedonic valence. Sounds were delivered *via* MRI-compatible earphones, and imaging was performed on a 1.5T scanner. Data were analyzed using voxel-based and ROI-based statistics in SPM12 with small volume correction (SVC). Due to hearing anomalies or MRI artefacts, only 5 pigs were included in the final analysis.

Functional MRI revealed that vocalizations activated regions of the auditory pathway and the left amygdala (pFWE at peak < 0.05 after SVC for all), with specific differences between positive and negative sounds. Clusters of activated voxels covering part of hippocampal areas, caudate nuclei and putamen were found with both positive and aversive vocal sounds. Limbic regions, including the amygdala and insula (p<0.05), as well as the right hippocampus after SVC (pFWE = 0.015) were uniquely engaged during the perception of negative conspecific vocalizations, indicating distinct processing based on emotional valence.

This study shows for the first time that piglets’ brain can process and differentiate emotional vocalizations from other pigs, even under general anesthesia. Positive and negative vocal sound playbacks activated distinct brain regions related to hearing, emotion and reward. These findings highlight pigs’ cognitive and emotional processing of vocal cues. This study is part of a wider research program aimed at developing the fMRI protocol with acoustic stimulation in juvenile pigs.

## 1. Introduction

In the realm of animal welfare science and affective ethology, vocalizations have emerged as a rich and non-invasive proxy to explore the emotional lives of animals. Among domesticated species and like many other social mammals, pigs (*Sus scrofa domesticus*) use vocalizations to communicate static (*i*.*e*. identity) and dynamic information (health, physiological status and emotions). Indeed, vocalizations are linked to the anatomy of the individuals, and so can encode descriptive information like the identity of the individual or its size; but they may also vary with the physiological status of the animals, and encode more dynamic information [1, 2]. Numerous recent studies have investigated how pigs express emotions through a complex and nuanced vocal repertoire (Denis et al., 2025), their meaning across contexts [3], and how conspecifics [4], humans [5] or even artificial intelligence systems [6] can interpret them.

Emotions in animals may be conceptualized according to their intensity (arousal) and valence (positive or negative) [7]. Pigs, like other mammals, modulate their vocal productions according to their emotional state along both axes. Tallet et al. [8] provided a foundational acoustic analysis of piglet vocalizations recorded across 11 situations, either positive or negative (*e*.*g*. castration, fighting, isolation, social reunion, nursing, *etc*.). Through quantitative characterization of 1,513 calls, the authors demonstrated that vocal expressions can be grouped into 2 to 5 call types, aligning with discrete emotional situations, and suggesting a nuanced and graded expression of affective states, as corroborated in wild boars by Garcia et al. [9]. Briefer et al. [10] later showed that domestic pig grunts differ significantly between positive and negative contexts in their spectral and temporal features, with grunts produced in a positive situation characterized by higher filter-related parameters (*i*.*e*. formants), a shorter duration, a lower fundamental frequency and a lower harmonicity compared to negative grunts. This was confirmed at a larger scale by Briefer et al. [6], who showed that with only 4 acoustic features, one is able to accurately categorize the calls into positively and negatively valenced. Similar findings were echoed by Friel et al. [11], who reported that grunts were shorter during positive cognitive bias trials, albeit without detecting broader acoustic shifts found in other studies. Importantly, not all vocalizations are uniformly impacted by emotional states. Linhart et al. [12] found that the effect of arousal on vocal parameters depends on the function of the vocalization type. In distress calls, higher arousal correlated with greater intensity and harmonicity, while in contact calls, the harmonic-to-noise ratio decreased with arousal, implying that different call types serve distinct communicative functions and may have evolved according to different evolutionary pressures. Being conditioned to be reunited with conspecifics or to a familiar human also shapes vocalizations’ structure, as spectrotemporal features of grunts varied according to the expected social partner [13], reflecting different expectations based on different valence and arousal states. The same authors showed different vocal responses of pigs interacting with a familiar human compared to a familiar inanimate object, suggesting a more reassuring and emotionally enriching effect of the human presence [14]. Thus, vocalizations are clear expressions of inner state.

Despite these strides, a major gap remains concerning the neural mechanisms underpinning emotionally charged vocalizations. Most of the studies focusing on the neural mechanisms of vocal communication in animals were performed in birds [15], but comparative studies do exist in birds, rodents and non-human primates [16]. While behavioral and acoustic studies have flourished, neuroscientific insights into pig vocalization are just beginning to emerge. For example, Leliveld et al. [4] tackled lateralization of vocal processing in pigs, showing that restraint calls induced a left head-turning bias, indicative of a right hemispheric dominance, while isolation calls were more processed by the left hemisphere. These patterns mirror human hemispheric specialization, left for linguistic, right for emotional. Palma et al. [17] argued that pig studies can illuminate speech and emotion evolution. Though, unlike primates or birds, pigs remain underused in cognitive neuroscience. To bridge this gap, implementation of neuroscientific techniques such as fMRI, fNIRS, EEG or portable biosensors could map brain activity during vocal events in emotion- and vocal-related regions (*e*.*g*. amygdala, periaqueductal gray, anterior cingulate cortex, auditory and motor cortex). Simultaneous recordings of vocal output, physiological arousal and brain activity would offer a holistic view of how emotion are encoded. To date, only a few studies have used pig models in hearing research for the development of new cochlear implants [18, 19], however, detailed brain responses to emotional calls using high-resolution imaging remain unexplored.

Functional brain imaging (*i*.*e*. blood-oxygen-level-dependent functional magnetic resonance imaging – BOLD fMRI – or nuclear imaging such as positron emission tomography – PET) has already been implemented in pig models and extensively used to investigate brain responses to sensory signals with different hedonic values, including odors, tastes and flavors [20 for review]. For example, our group demonstrated that the perception of flavors previously associated with positive or negative post-ingestive consequences modulated brain responses in the corticolimbic circuit, including the basal and thalamic nuclei, prefrontal cortex, cingulate cortex and temporal gyrus [21, 22]. Familiarity to an orexigenic functional feed ingredient was also found to modulate brain responses in reward and memory regions including the amygdala, insular cortex, prepyriform area and dorsal striatum [23]. More recent work using BOLD fMRI investigated the brain responses to the oral perception of sucrose or quinine [24], the olfactory perception of putatively orexigenic functional food ingredients [25], or the effects of a food ingredient on the brain responses to acute or chronic stressors [26, 27]. These examples demonstrate the feasibility of exploring hedonic processing and emotion perception in pigs under general anesthesia.

Building on this, our study aims to: i) implement BOLD fMRI to examine brain auditory responses in pigs, ii) compare brain responses to conspecific vocal signals of different emotional valence, iii) interpret these data in light of cognition, welfare, and translational neuroscience. We hypothesize that vocal signals with different emotional valence and meaning trigger different functional responses in brain structures involved in hearing processing, reward, learning and memory, with a particular focus on limbic areas. More precisely, we hypothesize positive calls will recruit reward regions and negative calls the amygdala, both of them also activating hippocampal regions.

## 2. Materials and methods

### 2.1. Animals

Experiments were conducted in accordance with the current ethical standards of the European Community (Directive 2010/63/EU), Agreement No. D3527532 and Authorization No. 26621-2020071709294199. The Regional Ethics Committee in Animal Experiment of Brittany has validated and approved the entire procedure described in this paper (project number 2020071709294199).

A total of eight 2-month-old 25-kg Piétrain × (Large White/Landrace) pigs were used in this study. There were 4 females and 4 males born at the INRAE experimental research station of Saint-Gilles (Pig Physiology and Phenotyping Experimental Facility, doi:10.15454/1.5573932732039927E12). Before imaging on the PRISM platform (Rennes), the pigs were temporarily (less than 24h) housed in individual pens (150 × 60 × 80 cm) and had free access to water. At least two animals were housed in the animal facility to permit visual and auditory contacts. The room was maintained at ∼ 24 °C with a natural 13:11 light–dark cycle. Criteria for inclusion in the final imaging analysis were defined *a priori* as: (i) normal bilateral hearing, defined by auditory brainstem response (ABR) thresholds ≤ 60 dB at 250 Hz, 3000 Hz and 8000 Hz; and (ii) absence of morphological ear abnormalities on anatomical T1-weighted MRI. Animals not meeting these criteria, or presenting major MRI artefacts, were excluded from the final fMRI analysis but not from the experimental procedures.

### 2.2. Anesthesia

For ABR and fMRI, pre-anesthesia was performed with an intramuscular injection of TILETAMINE/ZOLAZEPAM (15 mg/kg — Zoletil 50^®^ 25mg/ml, Virbac, Carros, France) in overnight-fasted animals. ISOFLURANE 5% inhalation (Isoflu-Vet 1000mg/g^®^ 250 mL, Dechra, Montigny-le-Bretonneux, France) was used to suppress the pharyngotracheal reflex). After intubation, anesthesia was maintained with 2% ISOFLURANE and mechanical ventilation allowed adjustment of respiratory frequency at 16 breaths/min with a tidal volume of 380 mL (Fabius, Draeger^®^, Germany). The end tidal volume was adapted in order to target etC02 between 4% and 4.5%. Heart rate was comprised between 100 and 150 beats per minute. The body temperature was recorded with intra-rectal thermal probe. Animals were covered with a blanket during imaging and tape was used to keep the eyes closed. Animals were euthanized at the end of the imaging session with intra-veinous injection of EMBUTRAMIDE / MÉBÉZONIUM / TÉTRACAÏNE (1mL/10kg, T61 50mL^®^, MSD, Beaucouze, France) without awakening from anesthesia.

### 2.3. Design of the study

Evoked ABR was realized on all pigs to ensure the integrity of their hearing, under general anesthesia. Then, anatomical MRI at 1.5 T was performed to visualize the brain and auditory pathways. Next, BOLD fMRI was performed with different pig vocalization with positive or negative emotional valence.

### 2.4. Auditory stimulation

Foam Caps (ER3-14 A, Echodia, France) were placed in the bony external ear canal of each ear. These caps were connected *via* a silicone tube to Insert Earphone (E-A-Rtone 3A®, 3M E-A-R^TM^). The silicone tube was 1.5 m long. This distance was adjusted beforehand in order to find the best compromise between a distance that was too close and would deactivate the earpieces because of the magnetic field of the MRI, and a distance that was too far and would lead to a loss of intensity of the acoustic stimulation signal. Quick-setting silicone (Instamold^®^, Interson Protac, France) was used within the pinna to attach the device and provide additional isolation from surrounding noise. The inserts were connected to the audiometers located outside the Faraday cage *via* speaker cables. To avoid any artifacts in the images due to parasitic signals transmitted by the loudspeaker cables in the Faraday cage, low-pass filters have been inserted to allow only the electrical signal related to the acoustic stimulation to pass through [28]. (***Supplementary Figure 1***).

The sound was generated by 2 audiometers (Acoustic Analyzer AA30, Starkey®) (Clinical audiometer AC30, Interacousitcs®) one for each ear. We used vocalizations from conspecifics from Tallet et al. [5] already used to test human perception with different intensity, hedonic valence and/or ecological meaning.

The vocal sound stimulation paradigm included five conditions (4 recordings and 1 silent control): Piglets being castrated [CAS] (high intensity, negative valence), piglets being isolated from their conspecifics [ISO] (medium intensity, negative valence), piglets reuniting with their conspecifics [REU] (medium intensity, positive valence), piglets after being nursed by their mother [NUR] (low intensity, positive valence) and silent during 15 seconds, repeated 16 times. All stimulus and silent blocks were presented in a pseudo-randomized order, including the silent condition. Consequently, identical conditions, including silence, could occasionally occur in consecutive blocks. The situations of recordings were fully described in Tallet et al. [5].

To further characterize the acoustic properties of the vocal stimuli, spectrotemporal representations and speechmaps were generated. Speechmaps illustrate the distribution of acoustic energy across frequency and time and allow visualization of differences in spectral content between emotional vocalizations (**Figure 1**).

**Figure 1.**
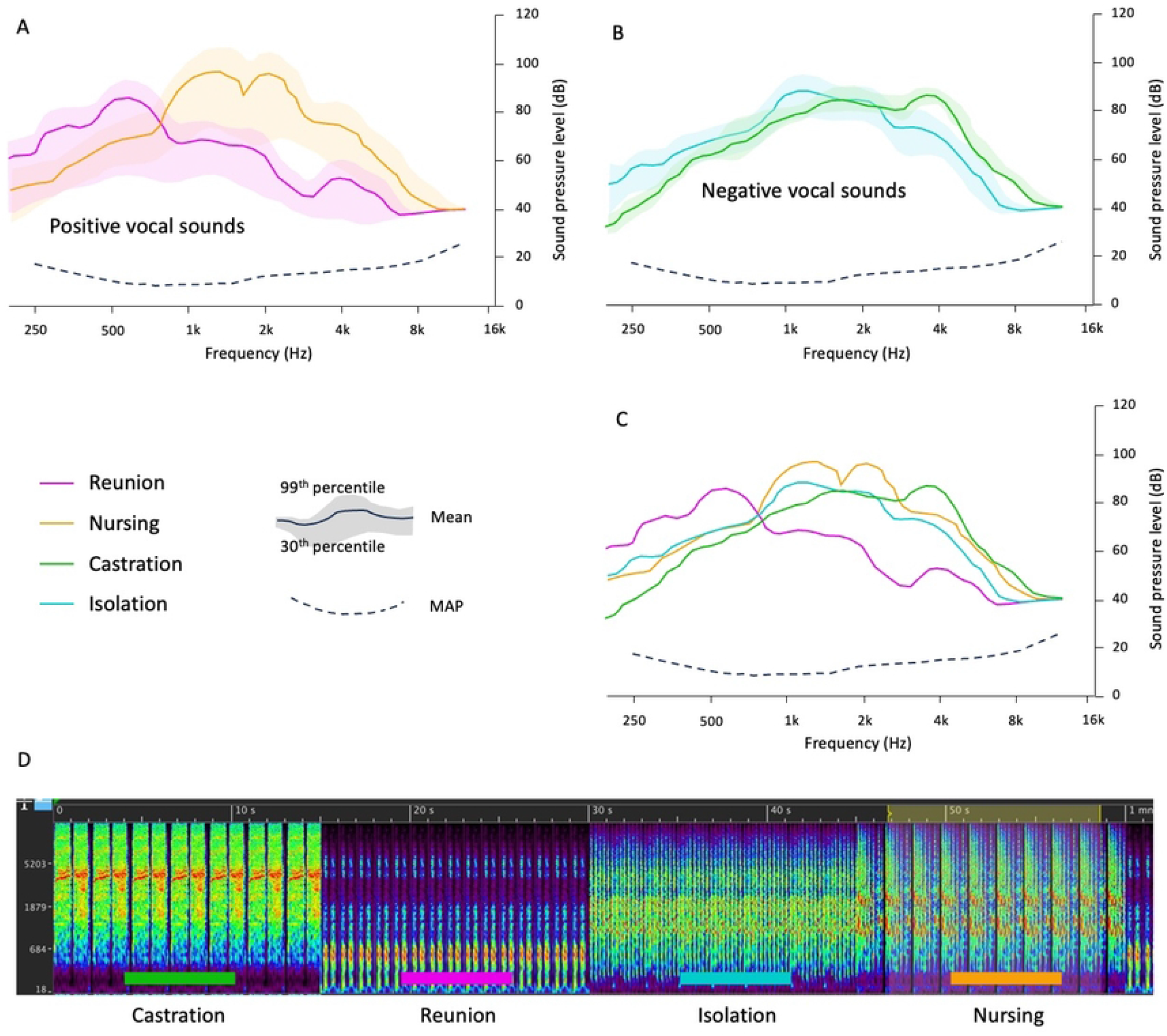
Acoustic characteristics of the vocal sounds used as auditory stimulations during BOLD fMRI brain imaging in pigs. **A.** Speechmaps (with mean sound pressure levels, 30^th^ and 99^th^ percentiles) of the two positive vocalizations, *i*.*e*. reunion (pink) and nursing (orange). **B**. Speechmaps (with mean sound pressure levels, 30^th^ and 99^th^ percentiles) of the two negative vocalizations, *i*.*e*. castration (green) and isolation (blue). **C**. Speechmaps (with mean sound pressure levels only) of the four vocalizations. **D**. Sonagram cumulating 15 seconds of each vocalization type. MAP: Minimum Audible Pressure, which is the sound pressure level at the eardrum corresponding to the threshold for normal hearing.

### 2.5. *In vivo* acoustic measurements

The sound conditions under which MRI is performed (noisy environment) were experimentally reproduced on an anaesthetized pig to verify acoustic detectability of stimuli relative to MRI acoustic noise. This measurement was performed by a Verifit2 device (Audioscan®, Ontario, Canada). The MRI noise was previously recorded and its spectral and intensity analysis was performed. This noise was simulated by a loudspeaker near the pig’s ear at an intensity of 120 dB SPL. Measurements of the sound band with the *in vivo* probe were carried out under different conditions: (i) at the entrance to the pinna, (ii) in the EAC (External Auditory Canal) without insert to measure the natural attenuation of the pinna and its cartilaginous reliefs (iii) in the EAC with insert to measure the level of natural attenuation of the pinna and the acoustic stimulation system (iv) with paste silicon in the EAC and pinna.

### 2.6 MRI image acquisition

Imaging was performed on a 1.5-T magnet (Siemens Avanto, Erlangen, Germany) at the Rennes Platform for Multimodal Imaging and Spectroscopy (PRISM). Acquisitions were performed using a combination of a Spine coil and a Flexible Body coil (Tim coil, Siemens^®^, Germany) placed on the head of the pig.

#### T1 weighted anatomical image acquisition

A magnetization-prepared rapid 3D gradient-echo (MP-RAGE) sequence was adapted for young conventional pig anatomy (voxel size: 1.2 × 1.2 × 1.2 mm^3^, FOV = 229 × 229 × 192 mm^3^, matrix = 192 × 192 × 160, number of averages (NA) = 2, repetition time (TR) = 2400ms, echo time (TE) = 3.62ms, flip angle (FA) = 8°, inversion time (IT) = 854ms, acquisition duration 15 minutes).

#### BOLD signal acquisition

A 2D echo planar imaging (EPI) sequence was adapted for pig head geometry (32 slices in ascending order, TR/TE: 2500/40ms, FA: 90°, voxel size: 2.5 × 2.5 × 2.5 mm^3^, FOV = 192 × 192 mm, matrix = 64 × 64). The EPI imaging time was 20 min 30 (497 volumes × 2.5 seconds/volume). The first four acquired volumes were excluded for the data analysis, meaning that no stimulation was performed during this period.

### 2.7 Data analysis and statistical image analysis

Data analysis was performed with SPM12 (version 6906, Wellcome Dept. of Cognitive Neurology, London, UK). Pre-processing of the BOLD images was carried out prior to the statistical analysis. Slice timing correction was performed using the 16th slice as reference. Realignement was based on the first volume, and a criterion of one voxel (scan-to-scan) was used to exclude volumes with too much motion. Spatial coregistration between BOLD images and the anatomical reference was calculated using the first EPI volume and the resulting transformation was applied to the rest of the sequence. Normalization was conducted with a 3-class tissue probability map (TPM) coregistered with the pig brain atlas (Saikali *et al*. 2010) and applied to the BOLD images. Then, images were smoothed with a Gaussian kernel of 4 mm, to improve the signal-to-noise ratio. Due to limitations related to the size of the pig brain and the effect of anesthesia on brain activity, we used a non-standard statistical analysis with regards to human statistical standards usually considering statistical significance at a cluster level with p-value < 0.05 under FDR correction, see details below. Anesthesia has indeed an effect on brain vascular reactivity. It has been shown that blood flow reactivity to CO2 challenge is reduced under isoflurane [29]. In dogs, BOLD signal variation under isoflurane anesthesia (about 1% signal variation), is detected as lower than human in awake state (about 5%) [30]. This reduced BOLD signal response prevents the use of human statistical standards, that are based on awake human brain. Indeed, lower variation on BOLD signal in anesthetized animals will consequently impair the detectability of human-standard statistics. In animals, only intracranial stimulation has shown sufficient level of stimulation allowing the use of human statistical standard, as discussed in Coquery *et al*. [24]. Also, the configuration used in this experiment is unusual and differs from that used in human studies. Unfortunately, it was not possible to use a standard multi-element head coil due to anatomical constraints. We therefore used a flexible body-coil combined with a spine-coil with 6 elements for each coil. Furthermore, the bones of the pig skull are much thicker than those of humans, which ‘mechanically distances’ the brain from these antennas. Of course, this configuration is not as effective as a standard coil (32 or 64 elements), but this type of configuration with surface coils combining a total of 12 elements has good sensitivity and offers the advantage of being closer to the animal’s skull. Although this does not entirely compensate for the loss of SNR compared to a ‘clinical’ configuration, this configuration still allows for a decent SNR to be obtained, provided that a compromise is made on statistical requirements.

#### Voxel-based statistic

First-level (within-individual contrast) and second-level (within-group contrast) t-test statistics were assessed for each voxels comparison with a threshold set at p < 0.05.

#### *SVC-based statistics* (Small Volume Correction)

Anatomical regions of interest (ROIs) from the Saikali pig brain atlas (Saikali *et al*. 2010) were used for SVC-based statistics with a p-value family-wise error (FWE) corrected at peak. We also performed a ROI analysis with multiple ROI correction statistic using MarsBar as previously described [24]. For the SVC and ROI analysis, fourteen ROIs corresponding to seven bilateral brain structures were selected based on *a priori* hypothesis (i) auditory sensory brain regions: Inferior colliculi (IC, 100 voxels each side), geniculate nuclei (GN, 120 voxels each side) and auditory cortex (AC, 1075 voxels each side); (ii) Motivational and reward brain regions: Caudate nucleus (CN, 615 voxels each side) and putamen (Put, 531 voxels each side); (iii) Limbic and associative learning brain regions: Amygdala (AMY, 230 voxels each side) and hippocampus (HIP, 440 voxels each side). For voxel-based statistics and SVC-based statistics, no suprathreshold voxels were detected with FDR (False Discovery Rate) correction at p < 0.05. Due to technical problems during functional acquisition, *i*.*e*., artefacts, and hearing loss, only five animals were used for analysis. The experimental setup, the acquisition and the statistical analysis has been previously validated in a previous study (Coquery *et al*. 2018).

## 3. Results

### 3.1. Pigs

The juvenile pigs were on average 86 days old (minimum 68 days / maximum 105 days). Their average live body weight was 28 kg (min 23.6 kg / max 34 kg). Anesthetic induction was performed on average with 2.25 mL of TILETAMINE/ZOLAZEPAM (min 1.5 mL / max 3 mL) combined with 5% ISOFLURANE. The average duration of anesthesia was 5 hours and 30 minutes (min 4 hours 10 minutes / max 8 hours). Physiological parameters during anesthesia averaged 134 beats per minute for heart rate (min 100 / max 167), 98% for oxygen saturation (min 96% / max 99%), 33.9°C for temperature (min 33.1°C / max 37.1°C), 4.1% for EtCO2 (min 3.9 / max 4.5) and 1.8 (min 0.9 / max 2.2) and 1.6 (min 0.9 / max 2) for the fraction of inspired and exhaled ISOFLURANE respectively. There was no complication during anesthesia.

Three pigs were excluded on the basis of anatomical and/or functional MRI, meaning that data from 5 pigs were included in the statistical analyses. The anatomical MRI, used to interpret the fMRI acquisitions, showed a normal anatomy for most of the animals (n= 6). One pig presented a unilateral deafness corresponding to a unilateral increase in audiometric thresholds which may correspond to a conductive hearing loss due to middle ear damage. The analysis of the anatomical images of this pig revealed an abnormal signal of the tympanic cavity which seemed “filled” without being able to determine the nature of this anatomical anomaly. It was decided to exclude this pig from the final analysis of the binaural acoustic stimulation model. This same anomaly was found in a second pig, for which artefacts in the BOLD signal were also found in addition to an identical deafness. This same pig and a third one presented significant artefacts in ROIs such as the temporal cortex, making the results uninterpretable.

### 3.2. *In vivo* acoustic measurements

The maximum attenuation of the noise generated by the MRI with insert and silicone in the pinna was 40-45dB. This estimation was probably a low estimate because in real experimental conditions: (i) better hermetic of the measurement system due to the absence of leakage from the *in vivo* measurement probe (ii) insert closer was closer to the eardrum (iii) real MRI noise level probably lower (high range taken for manipulation, about 115dB). All this consideration, enabled the signal-to-noise ratio to be estimated at 15dB.

### 3.3. Functional MRI: Brain responses to vocal acoustic stimulations

#### Brain responses to vocal acoustic stimulation

With all four vocal stimulations combined, we found clusters of activated voxels covering part of the IC, the GN and the AC (**Figure 2A** and **Table 1**). After ROI-based analysis, we could detect a corrected increase of the BOLD signal in the left IC (p-cor=0.011). After SVC-based statistic, we could detect a corrected increase of BOLD signal in the right GN (pFWE at peak =0.042), the left AC (pFWE at peak=0.027), the left IC (pFWE at peak=0.023), the left AMY (pFWE at peak=0.035).

**Table 1.** Whole-brain analysis of the brain BOLD fMRI responses of pigs to vocal sound stimulation with an uncorrected voxel-level p_value_ set at 0.05. Only the peaks at p<0.001 are presented here. The complete table is provided in Supplementary data. Cluster and peak statistics are presented with uncorrected, False-Wise Error Rate (FWER) and False Discovery Rate (FDR) corrected p values. k_E_ represents the cluster extent in voxels, T the t-value and Z_E_ the corresponding z-value. The coordinates in the dorsal-ventral position related to the posterior commissure (in mm) are indicated as x, y, and z. ROI is indicated when the peak is inside a region of the pig brain atlas.

| ROI | cluster-level |  |  | peak-level |  |  |  |  |  |  |
| --- | --- | --- | --- | --- | --- | --- | --- | --- | --- | --- |
| | $p_{\text{FWER-corr}}$ | $p_{\text{FDR-corr}}$ | $k_E$ | $p_{\text{uncorr}}$ | $p_{\text{FWER-corr}}$ | $p_{\text{FDR-corr}}$ | $T$ | $(z_E)$ | $p_{\text{uncorr}}$ | $x, y, z$ |
| Inferior cerebellar peduncle, L Substantia nigra,<br>L Amygdala, L Piriform cortex | 0.007 | 0.578 | 1724 | <0.001 | 0.342 | 1.030 | 30.09 | 4.49 | <0.001 | -7 -8 -7 |
| Crus 1 of the ansiform lobule | 1.000 | 0.935 | 35 | 0.487 | 1.000 | 1.030 | 18.04 | 4.03 | <0.001 | 6, 8, -27 |
| L Auditory Cortex L Insular cortex | 1.000 | 0.935 | 26 | 0.554 | 1.000 | 1.030 | 16.09 | 3.92 | <0.001 | -23, 13, 7 |
| R Lateral ventricle, R Caudate nucleus,<br>Corpus callosum | 1.000 | 0.935 | 32 | 0.508 | 1.000 | 1.030 | 12.46 | 3.67 | <0.001 | 2, 5, 22 |
| R Mediodorsal thalamic nucleus<br>L Mediodorsal thalamic nucleus | 1.000 | 0.935 | 21 | 0.599 | 1.000 | 1.030 | 10.61 | 3.51 | <0.001 | 1, 3, 6 |
| R Putamen | 1.000 | 0.935 | 52 | 0.392 | 1.000 | 1.030 | 9.95 | 3.44 | <0.001 | 7, -4, 28 |
| Medial cerebellar peduncle,<br>R Pontine nuclei | 1.000 | 0.935 | 43 | 0.438 | 1.000 | 1.030 | 9.55 | 3.40 | <0.001 | 7, -12, -9 |
| R Amygdala | 1.000 | 0.935 | 166 | 0.131 | 1.000 | 1.030 | 8.62 | 3.29 | <0.001 | 11, 0, 10 |
| R Primary Somatosensory Cortex | 1.000 | 0.935 | 10 | 0.732 | 1.000 | 1.030 | 8.31 | 3.25 | 0.001 | 11, 23, 17 |
| R Geniculate nuclei, R Perirhinal cortex,<br>R Parahippocampal cortex, R Anterior<br>entorhinal cortex | 0.999 | 0.935 | 245 | 0.072 | 1.000 | 1.030 | 7.48 | 3.14 | 0.001 | 8, -3, -3 |
| R Prepyriform area | 1.000 | 0.935 | 72 | 0.312 | 1.000 | 1.030 | 7.35 | 3.12 | 0.001 | 14, 1, 29 |
| R Inferior temporal gyrus, R Auditory<br>Cortex,<br>R Middle temporal gyrus | 1.000 | 0.935 | 26 | 0.554 | 1.000 | 1.030 | 7.30 | 3.11 | 0.001 | 25, 5, 1 |
| Crus 2 of the ansiform lobule | 1.000 | 0.935 | 11 | 0.717 | 1.000 | 1.030 | 7.09 | 3.08 | 0.001 | 15, -2, -11 |
| R Spinal trigeminal tract, R Spinal<br>trigeminal nucleus,<br>R Cuneate nucleus, R Hypoglossal<br>nucleus, L Hypoglossal nucleus | 1.000 | 0.935 | 199 | 0.101 | 1.000 | 1.030 | 6.95 | 3.06 | 0.001 | 8, -8, -21 |
| R Premotor Cortex, R Dorsal anterior<br>cingulate cortex,<br>L Lateral ventricle, L Putamen | 0.886 | 0.935 | 436 | 0.021 | 1.000 | 1.030 | 6.77 | 3.02 | 0.001 | 2, 15, 21 |
| L Superior temporal gyrus, L Middle<br>temporal gyrus,<br>L Pulvinar nuclei, Third ventricle, L<br>Subiculum | 1.000 | 0.935 | 116 | 0.202 | 1.000 | 1.030 | 6.47 | 2.97 | 0.001 | -23, 9, -4 |

**Figure 2.**
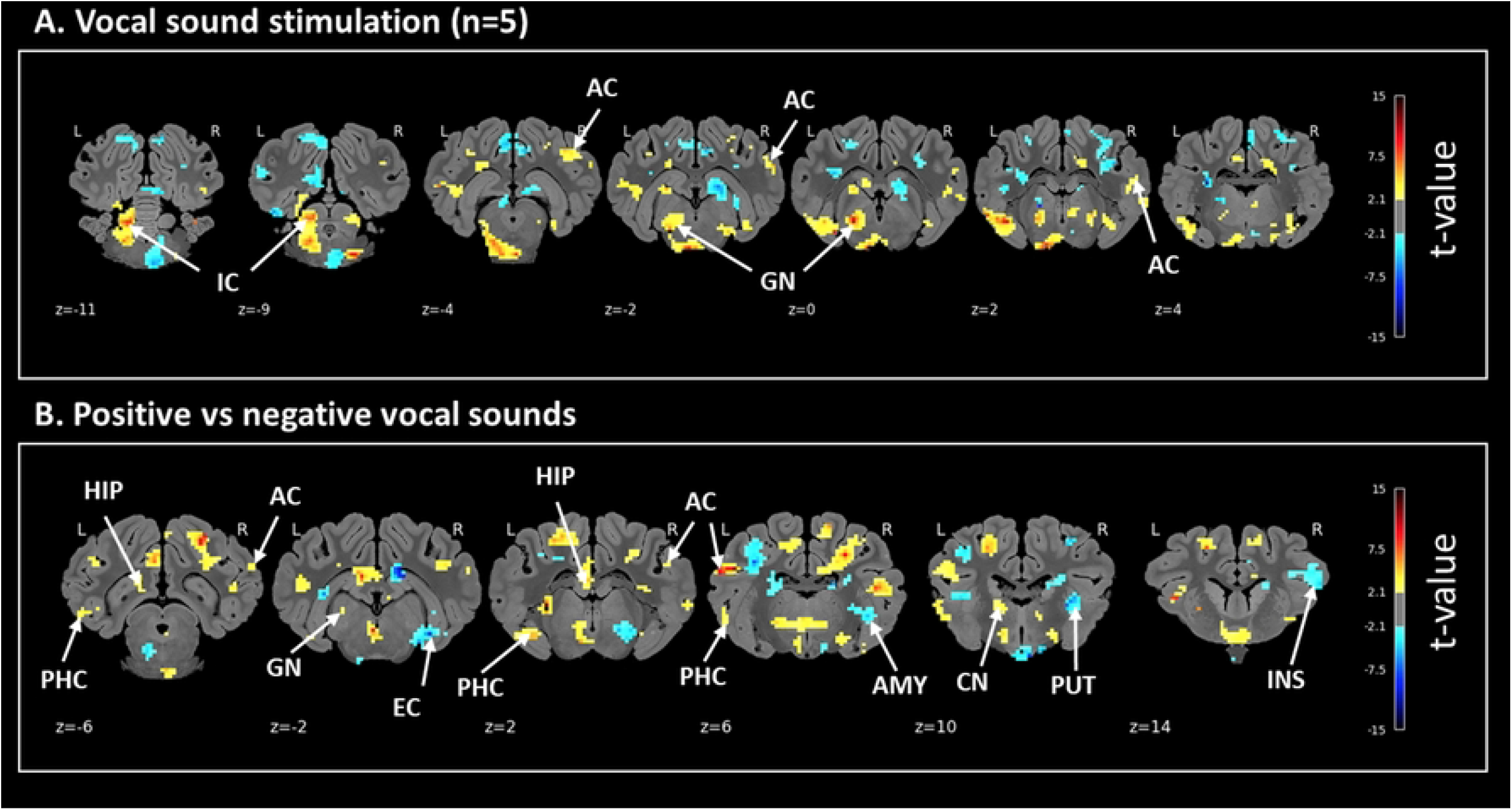
Maps of brain BOLD fMRI responses of pigs to (A) vocal sound stimulation and (B) positive versus negative vocal sounds stimulation. An uncorrected statistical voxel-level p_value_ = 0.05 was applied. The coordinates in the dorsal-ventral position related to the posterior commissure (in mm) are indicated below each slice level. Arrows indicate regions of interest in the Saikali atlas (AC: auditory cortex, AMY: amygdala, CN: caudate nucleus, EC: entorhinal cortex, GN: geniculate nuclei, HIP: hippocampus, IC: inferior colliculi, INS: insular cortex, PHC: parahippocampal cortex, PUT: putamen). R: right; L: left.

#### Variations in brain responses according to positive or negative vocal sound playbacks

With positive vocal sounds (*i*.*e*. REU and NUR) compared with negative vocal sounds, we found clusters of activated voxels covering part of the auditory pathways, *i*.*e*., the GN, the IC and the AC (**Figure 2B** and **Table 2**). With negative vocal sounds (CAS and ISO) compared with positive vocal sound, we found clusters of activated voxels covering part of the AC and the GN (**Figure 2B** and **Table 3**).

**Table 2.** Whole-brain analysis of the brain BOLD fMRI responses of pigs to positive *versus* negative vocal sounds stimulation with an uncorrected voxel-level p_value_ set at 0.05. Only the peaks at p<0.001 are presented here. The complete table is provided in Supplementary data. Cluster and peak statistics are presented with uncorrected, False-Wise Error Rate (FWER) and False Discovery Rate (FDR) corrected p values. k_E_ represents the cluster extent in voxels, T the t-value and Z_E_ the corresponding z-value. The coordinates in the dorsal-ventral position related to the posterior commissure (in mm) are indicated as x, y, and z. ROI is indicated when the peak is inside a region of the pig brain atlas.

| ROI | cluster-level |  |  | peak-level |  |  |  |  |  |  |
| --- | --- | --- | --- | --- | --- | --- | --- | --- | --- | --- |
| | $p_{\text{FWER-corr}}$ | $p_{\text{FDR-corr}}$ | $k_E$ | $p_{\text{uncorr}}$ | $p_{\text{FWER-corr}}$ | $p_{\text{FDR-corr}}$ | $T$ | $(z_E)$ | $p_{\text{uncorr}}$ | $x, y, z$ |
| L Primary Somatosensory Cortex | 1.000 | 0.935 | 104 | 0.224 | 0.258 | 1.031 | 32.30 | 4.55 | <0.001 | -12, 21, 19 |
| L Reticular thalamic nucleus, L Pulvinar nuclei, Periaqueductal gray, Cerebral aqueduct Sylvius, Third ventricle, R Ventral anterior thalamic nucleus, R Reticular thalamic nucleus | 0.405 | 0.347 | 695 | 0.005 | 0.521 | 1.031 | 27.07 | 4.40 | <0.001 | -12, 4, 2 |
| R Primary Somatosensory Cortex, R Dorsolateral prefrontal cortex, R Insular cortex | 0.998 | 0.935 | 270 | 0.059 | 1.000 | 1.031 | 20.37 | 4.14 | <0.001 | 9, 9, 38 |
| L Prepyriform area, L Insular cortex | 1.000 | 0.935 | 11 | 0.716 | 1.000 | 1.031 | 19.81 | 4.12 | <0.001 | -21, 8, 15 |
| R Dorsal anterior cingulate cortex Corpus callosum, R Ventral Anterior cingulate cortex, R Premotor Cortex, L Dorsolateral prefrontal cortex, R Anterior prefrontal cortex | 0.240 | 0.221 | 824 | 0.003 | 1.000 | 1.031 | 12.96 | 3.71 | <0.001 | 3, 12, 19 |
| L Somatosensory Association Cortex L Auditory Cortex, L Insular cortex | 1.000 | 0.935 | 117 | 0.199 | 1.000 | 1.031 | 11.89 | 3.63 | <0.001 | -19, 15, 6 |
| L Dorsal posterior cingulate cortex, L Lateral ventricle Corpus callosum, Fornix, L Hippocampus | 0.990 | 0.935 | 314 | 0.044 | 1.000 | 1.031 | 11.31 | 3.58 | <0.001 | -1, 18, -7 |
| L Fusiform gyrus L Parahippocampal cortex, Crus 2 of the ansiform lobule | 1.000 | 0.935 | 44 | 0.431 | 1.000 | 1.031 | 10.50 | 3.50 | <0.001 | -13, 4, -13 |
| R Somatosensory Association Cortex R Secondary visual cortex, R Primary visual cortex, R Dorsal posterior cingulate cortex Corpus callosum | 0.998 | 0.935 | 264 | 0.062 | 1.000 | 1.031 | 10.33 | 3.48 | <0.001 | 11, 20, 4 |
| L Primary visual cortex, L Secondary visual cortex, L Somatosensory Association Cortex, L Primary Motor Cortex L Somatosensory Association Cortex, L Primary Somatosensory Cortex | 0.992 | 0.935 | 308 | 0.046 | 1.000 | 1.031 | 9.82 | 3.43 | <0.001 | -8, 24, 3 |
| L Claustrum, L Prepyriform area, L Insular cortex | 1.000 | 0.935 | 24 | 0.570 | 1.000 | 1.031 | 9.73 | 3.42 | <0.001 | -16, 7, 13 |
| R Primary visual cortex R Secondary visual cortex | 1.000 | 0.935 | 173 | 0.123 | 1.000 | 1.031 | 9.68 | 3.42 | <0.001 | 10, 23, -6 |
| R Inferior temporal gyrus R Auditory Cortex | 1.000 | 0.935 | 10 | 0.731 | 1.000 | 1.031 | 9.19 | 3.36 | <0.001 | 25, 5, 3 |
| Crus 2 of the ansiform lobule Cerebellar lobule VIIIB | 1.000 | 0.935 | 29 | 0.528 | 1.000 | 1.031 | 8.97 | 3.34 | <0.001 | -9, 0, -23 |
| L Spinal trigeminal nucleus | 1.000 | 0.935 | 161 | 0.135 | 1.000 | 1.031 | 8.69 | 3.30 | <0.001 | -5 -10 -18 |
| Crus 2 of the ansiform lobule | 1.000 | 0.935 | 25 | 0.561 | 1.000 | 1.031 | 8.34 | 3.26 | 0.001 | 8, 4, -23 |
| R Claustrum R Inferior temporal gyrus, R Lateral ventricle Fornix | 1.000 | 0.935 | 112 | 0.208 | 1.000 | 1.031 | 8.03 | 3.21 | 0.001 | 18, 10, 5 |
| L Parahippocampal cortex | 1.000 | 0.935 | 79 | 0.288 | 1.000 | 1.031 | 7.82 | 3.19 | 0.001 | -20, -4, 0 |
| L Primary Somatosensory Cortex | 1.000 | 0.935 | 86 | 0.268 | 1.000 | 1.031 | 7.57 | 3.15 | 0.001 | -13, 13, 32 |
| R Parahippocampal cortex R Fusiform gyrus, Cerebellar lobule VI, R Superior temporal gyrus, R Fusiform gyrus, Cerebellar lobule VI Cerebellar lobule V | 0.933 | 0.935 | 397 | 0.026 | 1.000 | 1.031 | 7.34 | 3.12 | 0.001 | 10, 8, -15 |
| R Pontine nuclei, R Reticulotegmental nucleus of the pons, L Pontine nuclei, L Reticulotegmental nucleus of the pons | 1.000 | 0.935 | 44 | 0.431 | 1.000 | 1.031 | 7.08 | 3.08 | 0.001 | 0, -13, -5 |

**Table 3.**
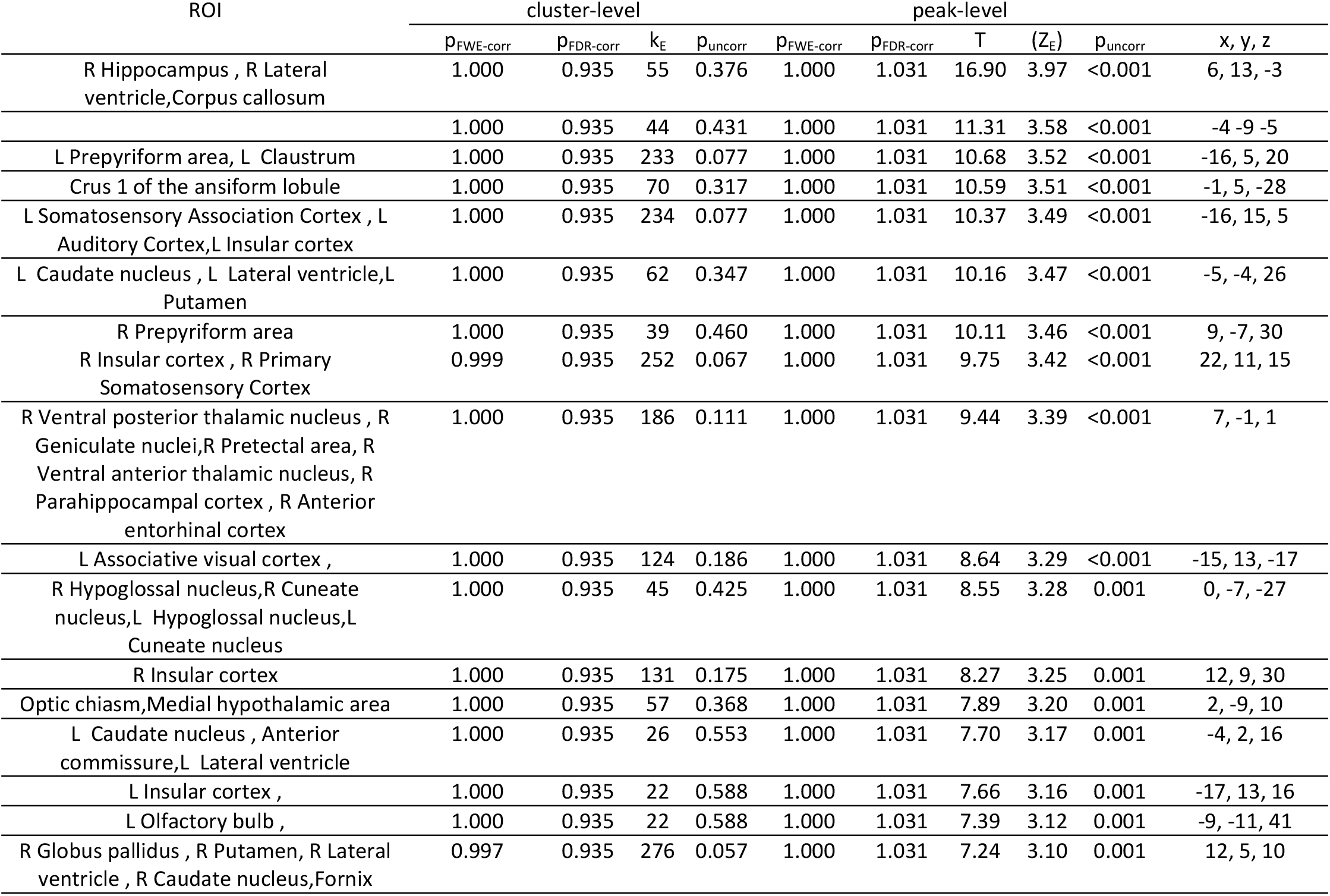
Whole-brain analysis of the brain BOLD fMRI responses of pigs to negative *versus* positive vocal sounds stimulation with an uncorrected voxel-level p_value_ set at 0.05. Only the peaks at p<0.001 are presented here. The complete table is provided in Supplementary data. Cluster and peak statistics are presented with uncorrected, False-Wise Error Rate (FWER) and False Discovery Rate (FDR) corrected p values. k_E_ represents the cluster extent in voxels, T the t-value and Z_E_ the corresponding z-value. The coordinates in the dorsal-ventral position related to the posterior commissure (in mm) are indicated as x, y, and z. ROI is indicated when the peak is inside a region of the pig brain atlas.

We found clusters of activated voxels covering part of the hippocampus (HIP), para-hippocampal cortex (PHC), caudate nuclei (CN) and putamen (PUT) with both positive and aversive vocal sounds. We found clusters of activated voxels covering part of other limbic structures such as the amygdala (AMY) and insular cortex (INS) only with negative vocal sound stimulation. After SVC-based statistic, we could detect a corrected increase of BOLD signal in the right HIP (pFWE at peak = 0.015) with negative *vs*. positive vocal sound stimulation.

#### Data sharing

Raw fMRI, anatomical and pre-processed data as well as activation maps are shared as open access data. DOI: https://doi.org/10.57745/NCR6XY

## 4. Discussion

In this study, we managed for the first time to describe the BOLD responses of piglet brains to conspecifics vocal sounds with different emotional valence. The first objective was to demonstrate that the brain of pigs subjected to general anesthesia could still respond to acoustic stimulation (*i*.*e*. vocalization playbacks), as we already showed for visual and olfactogustatory signals. Hence, we demonstrated that vocal sound perception activated the hearing neural pathway. Even more interesting is the fact that vocalizations emitted during a positive situation (reunion with conspecifics, after nursing by the mother) or a negative situation (castration, social isolation) led to different BOLD responses in brain structures involved in reward and learning, but also emotion perception and memory. This implies that different cognitive processes are triggered depending on the hedonic valence of vocal sounds from conspecifics and previous individual experience, and that these processes are still functional even under general anesthesia. This opens the way to further developments and studies in the context of fundamental and translational research in emotion communication and hearing sciences. For example, our pig model could be used to evaluate the emotional outcomes of particular auditory stimulations with the aim to improve animal welfare, or in the scope of preclinical research to develop new hearing therapies such as cochlear stimulation.

In previous studies, we have demonstrated in pigs that general anesthesia, when precisely monitored and controlled, does not prevent the animal’s brain from perceiving and integrating various types of sensory stimuli such as visual, olfactory, gustatory, olfactogustatory or even visceral (*e*.*g*. from the duodenum) signals [20]. We even showed that brain integration differs according to previous individual experience with these stimuli, implying that learning and memory processes (*i*.*e*. interpretation, even if it is unconscious) are still possible even under general anesthesia. Here, we showed that anesthetized pigs exposed to conspecifics’ vocalizations presented a significant activation of the brain auditory pathway, including the inferior colliculi (IC), geniculate nuclei (GN) and auditory cortex (AC). Other groups demonstrated that these brain structures are involved in functional plasticity in response to an acoustically enriched environment in rats [31], or in coding the temporal and spectral features of communication calls in the guinea pig [32]. In dogs, Bach *et al*. [33, 34] also showed activation of the caudal colliculi and medial geniculate nuclei in response to simple random noise stimuli. In addition to these activations of the auditory pathway, an activated area was even detected in the rostral aspect of the left caudate nucleus [34].

Beyond the mere demonstration of the auditory pathway activation in response to conspecifics vocalizations, our main goal was to investigate whether the cognitive and hedonic treatment of hearing conspecifics calls might differ according to their meaning and emission context. We hypothesized that positive vocal sounds would induce greater activity in the reward nuclei (including the dorsal striatum), in relation with the possible recall of pleasurable social interactions or emotional mirror processes (empathy), and that negative vocal sounds would induce greater activity in the amygdala, which is known to be involved in the treatment of fear, aversive learning and avoidance of dangerous situations [35-37]. In the human, the striatum releases dopamine when listening to pleasurable music while its activity codes the reward value of musical excerpts [38], and dorsal striatal activity is also detected during sympathetic concern [39]. The role of the amygdala in experiencing fear or triggering responses towards threatening or conflict situations has been extensively described in animal models [40], and a recent study showed that rats respond to aversive emotional arousal of human handlers with the activation of the basolateral and central amygdala [41]. Our first hypothesis was not validated since we found clusters of activated voxels covering part of the dorsal striatum (PUT and CN) with both negative and positive vocal sounds. This is surprising but it is important to remember that the dorsal striatum, in addition to being involved in pleasure and motivation, is also critical in controlling motor, procedural, and reinforcement-based behaviors. In the past, our group has demonstrated that the putamen and caudate nucleus activity could be modulated by both positive and negative sensory signals [21, 24], since functional subdivisions and lateralized responses do exist. Our second hypothesis was validated because we found clusters of activated voxels covering part of other limbic structures such as the amygdala (AMY) and insular cortex (INS) only with negative vocal sound stimulation, suggesting that the animals’ brain might have perceived the threatening, suffering or alert dimension of these calls. Interestingly, the GN and AC were differently activated with positive and negative vocal sounds, indicating that part of the auditory pathway is involved in the treatment of the emotional valence contained in the temporal and spectral features of these sounds, and responds preferentially to a positive hedonic valence. Such a role of the auditory cortex in emotion processing has already been extensively described, notably in the human [42], but the demonstration in the pig is new, even though not surprising. The activation of hippocampal structures (HIP, PHC) with both positive and negative vocal sounds is interesting in terms of memory processes and recall of previous experiences, and this effect was only significant after small volume correction (SVC) in the right HIP with negative *vs*. positive vocal sound stimulation. Many studies investigated the role of HIP in auditory fear conditioning in rodents, showing that hippocampus may have a general role for consolidation of remote associative memory through reactivation of memory trace [43] and that its connection with amygdala might permit the transmission of social information to synchronize threat response [44].

This princeps exploratory study has several limitations. First, the final number of animals is low, which limits the power of the brain imaging analyses and justified the adoption of less stringent statistical thresholds as compared with usual recommendations. Second, since we cannot dissociate the valence and intensity of the vocal sounds used to stimulate the animals during BOLD fMRI, we cannot exclude that part of the effects observed were attributable to intensity rather than emotional valence. Further studies should be performed with more subjects and call types to disentangle the respective roles of intensity, hedonic value and meaning in brain activations. Third, even though we published many articles in the pig model, showing that emotionally positive or negative olfactory and/or gustatory stimulations trigger different brain responses in anesthetized animals according to their individual experiences [20], these responses are probably very different from those we might obtain in awake animals, and the same reservations are necessary in the scope of the present study. When we talk about different brain integration and interpretation according to the emotional valence of vocal sounds, we refer to different activations of the limbic pathway between positive and negative vocal sounds for example, without extrapolating to any conscious process or behavioral response to these sounds. Finally, it was quite surprising to find 2 pigs out of 8, especially at such a young age, with an anatomical anomaly of the tympanic cavity, detected with MRI, along with a unilateral hearing loss, detected by increase of the audiometric thresholds during ABR. Whether such an anomaly was the consequence of a cholesteatoma, tumor or congenital malformation is not known, but this raises the question of the prevalence and welfare consequences of such hearing problems in animal husbandry, which remain undetected without specific testing. This also requires a particular vigilance on the subject exclusion ratio for further studies on auditory abilities in pigs.

As Palma et al. [17] noted, expanding hearing research beyond traditional models such as songbirds and primates is essential for reconstructing the broader phylogenetic roots of speech and emotion. More experimental studies involving neuropharmacological manipulations or brain stimulation could test causal relationship between brain activity and vocal emotion expression, and the pig model represents a real asset in this scope. Longitudinal developmental studies could also explore how these neural systems mature and whether domestication has altered them, as suggested by behavioral differences between domestic pigs and wild boar in Maigrot et al. [45]. Functional brain imaging studies in pigs could also improve our understanding of the neural coding of emotions through the temporal and spectral characteristics of vocal sounds, therefore paving the way towards better automated solutions using artificial intelligence for understanding the animals’ emotional status and needs in husbandry systems. Finally, our princeps fMRI study on the exploration of auditory and hearing-related cognitive abilities in pigs opens new possibilities in using this animal model for investigating new technological and methodological approaches for cochlear stimulation and implant design [18, 19]. This is notably our current goal since cochlear stimulation has been implemented in piglets with disabled hearing by our group, with promising results to come on brain responses to electrical, acoustic and bimodal stimulation.

In conclusion, our princeps exploratory brain imaging study suggested that even under general anesthesia, the pig’s brain can still hear, process and interpret vocal sound playbacks with different hedonic valence, highlighting some neural processes that underlie the emotional dimension of vocal communication in pigs. Such a demonstration opens the way to fundamental research in ethology but also to translational research in animal welfare and medical hearing sciences to offer new technologies in hearing prosthesis and cochlear implantation.

## 5. Acknowledgments

We would like to thank Carole Guérin for selecting the representative pig vocalization recordings that were used in this brain imaging study. We thank PRISM core facility (Biogenouest, France Life Imaging, Univ Rennes, Univ Angers, INRAE, CNRS, France) and Stephane Quellec for the access to the MRI machine, staff from animal facility UE3P of INRAE for their technical support, Julien Georges and Alain Chauvin, as well as Régis Janvier (NuMeCan, INRAE 1341 Univ Rennes, France) for their help during the animal experimentations. We also thank Stephane Laurent, audioprothesist, for his help on acoustic installation.

---

**Supplementary Fig 1.**
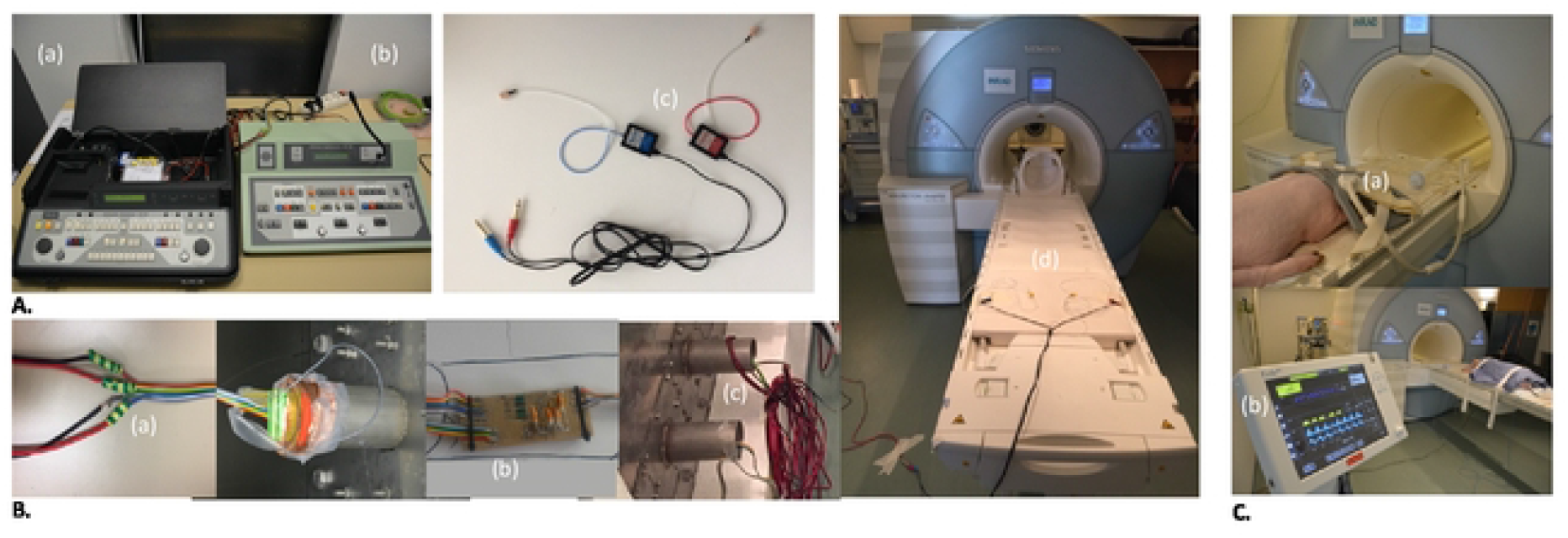
Pictures of the different installations. **(A)** Acoustic stimulation system: a) Acoustic Analyzer AA30 b) Clinical audiometer AC30 c) earphones with insert **(B)** Installation in the MRI room: a) soldering of the speaker cables to the filter outdoor the room b) frequency filter c) speaker cable inside the room d) MRI examination table with taped acoustic stimulation system **(C)** Anaesthetized pig: a) body coil on the head b) anesthesia monitor.

## References

1. Aubin T, Mathevon N. Coding strategies in vertebrate acoustic communication. Aubin T, Mathevon N, editors: Springer Cham; 2020.

2. Briefer EF, Padilla de la Torre M, McElligott AG. Mother goats do not forget their kids’ calls. Proc Biol Sci. 2012;279(1743):3749–55. Epub 20120620. doi: 10.1098/rspb.2012.0986. PubMed PMID: 22719031; PubMed Central PMCID: PMCPMC3415910.

3. Coutant M, Villain AS, Briefer EF. A scoping review of the use of bioacoustics to assess various components of farm animal welfare. Applied Animal Behaviour Science. 2024;275. doi: 10.1016/j.applanim.2024.106286.

4. Leliveld LMC, Dupjan S, Tuchscherer A, Puppe B. Hemispheric Specialization for Processing the Communicative and Emotional Content of Vocal Communication in a Social Mammal, the Domestic Pig. Front Behav Neurosci. 2020;14:596758. Epub 20201120. doi: 10.3389/fnbeh.2020.596758. PubMed PMID: 33328923; PubMed Central PMCID: PMCPMC7714956.

5. Tallet C, Spinka M, Maruscakova I, Simecek P. Human perception of vocalizations of domestic piglets and modulation by experience with domestic pigs (Sus scrofa). J Comp Psychol. 2010;124(1):81–91. doi: 10.1037/a0017354. PubMed PMID: 20175599.

6. Briefer EF, Sypherd CC, Linhart P, Leliveld LMC, Padilla de la Torre M, Read ER, et al. Classification of pig calls produced from birth to slaughter according to their emotional valence and context of production. Sci Rep. 2022;12(1):3409. Epub 20220307. doi: 10.1038/s41598-022-07174-8. PubMed PMID: 35256620; PubMed Central PMCID: PMCPMC8901661.

7. Mendl M, Burman OH, Paul ES. An integrative and functional framework for the study of animal emotion and mood. Proc Biol Sci. 2010;277(1696):2895–904. Epub 20100804. doi: 10.1098/rspb.2010.0303. PubMed PMID: 20685706; PubMed Central PMCID: PMCPMC2982018.

8. Tallet C, Linhart P, Policht R, Hammerschmidt K, Simecek P, Kratinova P, et al. Encoding of situations in the vocal repertoire of piglets (Sus scrofa): a comparison of discrete and graded classifications. PLoS One. 2013;8(8):e71841. Epub 20130813. doi: 10.1371/journal.pone.0071841. PubMed PMID: 23967251; PubMed Central PMCID: PMCPMC3742501.

9. Garcia M, Gingras B, Bowling DL, Herbst CT, Boeckle M, Locatelli Y, et al. Structural Classification of Wild Boar (Sus scrofa) Vocalizations. Ethology. 2016;122(4):329–42. Epub 20160216. doi: 10.1111/eth.12472. PubMed PMID: 27065507; PubMed Central PMCID: PMCPMC4793927.

10. Briefer EF, Vizier E, Gygax L, Hillmann E. Expression of emotional valence in pig closed-mouth grunts: Involvement of both source-and filter-related parameters. J Acoust Soc Am. 2019;145(5):2895. doi: 10.1121/1.5100612. PubMed PMID: 31153321.

11. Friel M, Kunc HP, Griffin K, Asher L, Collins LM. Positive and negative contexts predict duration of pig vocalisations. Sci Rep. 2019;9(1):2062. Epub 20190214. doi: 10.1038/s41598-019-38514-w. PubMed PMID: 30765788; PubMed Central PMCID: PMCPMC6375976.

12. Linhart P, Ratcliffe VF, Reby D, Spinka M. Expression of Emotional Arousal in Two Different Piglet Call Types. PLoS One. 2015;10(8):e0135414. Epub 20150814. doi: 10.1371/journal.pone.0135414. PubMed PMID: 26274816; PubMed Central PMCID: PMCPMC4537126.

13. Villain AS, Hazard A, Danglot M, Guerin C, Boissy A, Tallet C. Piglets vocally express the anticipation of pseudo-social contexts in their grunts. Sci Rep. 2020;10(1):18496. Epub 20201028. doi: 10.1038/s41598-020-75378-x. PubMed PMID: 33116261; PubMed Central PMCID: PMCPMC7595114.

14. Villain AS, Lanthony M, Guerin C, Tallet C. Manipulable Object and Human Contact: Preference and Modulation of Emotional States in Weaned Pigs. Front Vet Sci. 2020;7:577433. Epub 20201127. doi: 10.3389/fvets.2020.577433. PubMed PMID: 33330698; PubMed Central PMCID: PMCPMC7728720.

15. Louder MIM, Lawson S, Lynch KS, Balakrishnan CN, Hauber ME. Neural mechanisms of auditory species recognition in birds. Biol Rev Camb Philos Soc. 2019;94(5):1619–35. Epub 20190507. doi: 10.1111/brv.12518. PubMed PMID: 31066222.

16. Nieder A, Mooney R. The neurobiology of innate, volitional and learned vocalizations in mammals and birds. Philos Trans R Soc Lond B Biol Sci. 2020;375(1789):20190054. Epub 20191118. doi: 10.1098/rstb.2019.0054. PubMed PMID: 31735150; PubMed Central PMCID: PMCPMC6895551.

17. Palma M, Khoshnevis M, Lion M, Zenga C, Kefs S, Fallegger F, et al. Chronic recording of cortical activity underlying vocalization in awake minipigs. J Neurosci Methods. 2022;366:109427. Epub 20211128. doi: 10.1016/j.jneumeth.2021.109427. PubMed PMID: 34852254.

18. Fallegger F, Trouillet A, Lacour SP. Subdural Soft Electrocorticography (ECoG) Array Implantation and Long-Term Cortical Recording in Minipigs. J Vis Exp. 2023;(193). Epub 20230331. doi: 10.3791/64997. PubMed PMID: 37067278.

19. Yildiz E, Gerlitz M, Gadenstaetter AJ, Landegger LD, Nieratschker M, Schum D, et al. Single-incision cochlear implantation and hearing evaluation in piglets and minipigs. Hear Res. 2022;426:108644. Epub 20221031. doi: 10.1016/j.heares.2022.108644. PubMed PMID: 36343533.

20. Val-Laillet D. Review: Impact of food, gut-brain signals and metabolic status on brain activity in the pig model: 10 years of nutrition research using in vivo brain imaging. Animal. 2019;13(11):2699–713. Epub 20190729. doi: 10.1017/S1751731119001745. PubMed PMID: 31354119.

21. Clouard C, Jouhanneau M, Meunier-Salaun MC, Malbert CH, Val-Laillet D. Exposures to conditioned flavours with different hedonic values induce contrasted behavioural and brain responses in pigs. PLoS One. 2012;7(5):e37968. Epub 20120525. doi: 10.1371/journal.pone.0037968. PubMed PMID: 22685528; PubMed Central PMCID: PMCPMC3368353.

22. Gaultier A, Meunier-Salaun MC, Malbert CH, Val-Laillet D. Flavour exposures after conditioned aversion or preference trigger different brain processes in anaesthetised pigs. Eur J Neurosci. 2011;34(9):1500–11. Epub 20111017. doi: 10.1111/j.1460-9568.2011.07848.x. PubMed PMID: 22004412.

23. Val-Laillet D, Meurice P, Clouard C. Familiarity to a Feed Additive Modulates Its Effects on Brain Responses in Reward and Memory Regions in the Pig Model. PLoS One. 2016;11(9):e0162660. Epub 20160909. doi: 10.1371/journal.pone.0162660. PubMed PMID: 27610625; PubMed Central PMCID: PMCPMC5017780.

24. Coquery N, Meurice P, Janvier R, Bobillier E, Quellec S, Fu M, et al. fMRI-Based Brain Responses to Quinine and Sucrose Gustatory Stimulation for Nutrition Research in the Minipig Model: A Proof-of-Concept Study. Front Behav Neurosci. 2018;12:151. Epub 20180724. doi: 10.3389/fnbeh.2018.00151. PubMed PMID: 30140206; PubMed Central PMCID: PMCPMC6094987.

25. Coquery N, Menneson S, Meurice P, Janvier R, Etienne P, Noirot V, et al. fMRI-Based Brain Responses to Olfactory Stimulation with Two Putatively Orexigenic Functional Food Ingredients at Two Different Concentrations in the Pig Model. J Food Sci. 2019;84(9):2666–73. Epub 20190823. doi: 10.1111/1750-3841.14772. PubMed PMID: 31441517.

26. Briard E, Serrand Y, Dahirel P, Janvier R, Noirot V, Etienne P, et al. Exposure to a sensory functional ingredient in the pig model modulates the blood-oxygen-level dependent brain responses to food odor and acute stress during pharmacological MRI in the frontostriatal and limbic circuits. Front Nutr. 2023;10:1123162. Epub 20230228. doi: 10.3389/fnut.2023.1123162. PubMed PMID: 36925960; PubMed Central PMCID: PMCPMC10012862.

27. Menneson S, Serrand Y, Janvier R, Noirot V, Etienne P, Coquery N, et al. Regular exposure to a Citrus-based sensory functional food ingredient alleviates the BOLD brain responses to acute pharmacological stress in a pig model of psychosocial chronic stress. PLoS One. 2020;15(12):e0243893. Epub 20201228. doi: 10.1371/journal.pone.0243893. PubMed PMID: 33370353; PubMed Central PMCID: PMCPMC7769264.

28. Zhang X, Guerin S, Launay Y, Serrand Y, Coquery N, Val-Laillet D. Acute Effects of Different Electroacupuncture Point Combinations to Modulate the Gut-Brain Axis in the Minipig Model. Evid Based Complement Alternat Med. 2022;2022:4384693. Epub 20221020. doi: 10.1155/2022/4384693. PubMed PMID: 36310617; PubMed Central PMCID: PMCPMC9613379.

29. Munting LP, Derieppe MPP, Suidgeest E, Denis de Senneville B, Wells JA, van der Weerd L. Influence of different isoflurane anesthesia protocols on murine cerebral hemodynamics measured with pseudo-continuous arterial spin labeling. NMR Biomed. 2019;32(8):e4105. Epub 20190607. doi: 10.1002/nbm.4105. PubMed PMID: 31172591; PubMed Central PMCID: PMCPMC6772066.

30. Willis CKR, Quinn RP, McDonell WM, Gati J, Parent J, Nicolle D. Functional MRI as a tool to assess vision in dogs: the optimal anesthetic. Veterinary Ophthalmology 2001;4(4):243–53.

31. Svobodova Burianova J, Syka J. Postnatal exposure to an acoustically enriched environment alters the morphology of neurons in the adult rat auditory system. Brain Struct Funct. 2020;225(7):1979–95. Epub 20200625. doi: 10.1007/s00429-020-02104-8. PubMed PMID: 32588120.

32. Suta D, Popelar J, Syka J. Coding of communication calls in the subcortical and cortical structures of the auditory system. Physiol Res. 2008;57 Suppl 3:S149–S59. Epub 20080513. doi: 10.33549/physiolres.931608. PubMed PMID: 18481905.

33. Bach JP, Lupke M, Dziallas P, Wefstaedt P, Uppenkamp S, Seifert H, et al. Functional magnetic resonance imaging of the ascending stages of the auditory system in dogs. BMC Vet Res. 2013;9:210. Epub 20131016. doi: 10.1186/1746-6148-9-210. PubMed PMID: 24131784; PubMed Central PMCID: PMCPMC3854503.

34. Bach JP, Lupke M, Dziallas P, Wefstaedt P, Uppenkamp S, Seifert H, et al. Auditory functional magnetic resonance imaging in dogs--normalization and group analysis and the processing of pitch in the canine auditory pathways. BMC Vet Res. 2016;12:32. Epub 20160220. doi: 10.1186/s12917-016-0660-5. PubMed PMID: 26897016; PubMed Central PMCID: PMCPMC4761139.

35. LeDoux JE. Emotion circuits in the brain. Annu Rev Neurosci. 2000;23:155–84. doi: 10.1146/annurev.neuro.23.1.155. PubMed PMID: 10845062.

36. Olsson A, Phelps EA. Social learning of fear. Nat Neurosci. 2007;10(9):1095–102. doi: 10.1038/nn1968. PubMed PMID: 17726475.

37. Twining RC, Vantrease JE, Love S, Padival M, Rosenkranz JA. An intra-amygdala circuit specifically regulates social fear learning. Nat Neurosci. 2017;20(3):459–69. Epub 20170123. doi: 10.1038/nn.4481. PubMed PMID: 28114293; PubMed Central PMCID: PMCPMC5323274.

38. Zatorre RJ. Musical pleasure and reward: mechanisms and dysfunction. Ann N Y Acad Sci. 2015;1337:202–11. doi: 10.1111/nyas.12677. PubMed PMID: 25773636.

39. Kawamichi H, Tanabe HC, Takahashi HK, Sadato N. Activation of the reward system during sympathetic concern is mediated by two types of empathy in a familiarity-dependent manner. Soc Neurosci. 2013;8(1):90–100. Epub 20121119. doi: 10.1080/17470919.2012.744349. PubMed PMID: 23163952.

40. Leake J, Leidl DM, Lay BPP, Fam JP, Giles MC, Qureshi OA, et al. What is Learned Determines How Pavlovian Conditioned Fear is Consolidated in the Brain. J Neurosci. 2024;44(2). Epub 20240110. doi: 10.1523/JNEUROSCI.0513-23.2023. PubMed PMID: 37963767; PubMed Central PMCID: PMCPMC10860607.

41. Kazmierowska AM, Kostecki M, Szczepanik M, Nikolaev T, Hamed A, Michalowski JM, et al. Rats respond to aversive emotional arousal of human handlers with the activation of the basolateral and central amygdala. Proc Natl Acad Sci U S A. 2023;120(46):e2302655120. Epub 20231107. doi: 10.1073/pnas.2302655120. PubMed PMID: 37934822; PubMed Central PMCID: PMCPMC10655214.

42. Koelsch S, Skouras S, Lohmann G. The auditory cortex hosts network nodes influential for emotion processing: An fMRI study on music-evoked fear and joy. PLoS One. 2018;13(1):e0190057. Epub 20180131. doi: 10.1371/journal.pone.0190057. PubMed PMID: 29385142; PubMed Central PMCID: PMCPMC5791961.

43. Oh JP, Han JH. A critical role of hippocampus for formation of remote cued fear memory. Mol Brain. 2020;13(1):112. Epub 20200815. doi: 10.1186/s13041-020-00652-y. PubMed PMID: 32799906; PubMed Central PMCID: PMCPMC7429722.

44. Ito W, Palmer AJ, Morozov A. Social Synchronization of Conditioned Fear in Mice Requires Ventral Hippocampus Input to the Amygdala. Biol Psychiatry. 2023;93(4):322–30. Epub 20220804. doi: 10.1016/j.biopsych.2022.07.016. PubMed PMID: 36244803; PubMed Central PMCID: PMCPMC10069289.

45. Maigrot AL, Hillmann E, Briefer EF. Cross-species discrimination of vocal expression of emotional valence by Equidae and Suidae. BMC Biol. 2022;20(1):106. Epub 20220524. doi: 10.1186/s12915-022-01311-5. PubMed PMID: 35606806; PubMed Central PMCID: PMCPMC9128205.

